# Modeling cell-substrate de-adhesion dynamics under fluid shear

**DOI:** 10.1101/166371

**Authors:** Renu Maan, Garima Rani, Gautam I. Menon, Pramod A. Pullarkat

## Abstract

Changes in cell-substrate adhesion are believed to signal the onset of cancer metastasis, but such changes must be quantified against background levels of intrinsic heterogeneity between cells. Variations in cell-substrate adhesion strengths can be probed through biophysical measurements of cell detachment from substrates upon the application of an external force. Here, we investigate, theoretically and experimentally, the detachment of cells adhered to substrates when these cells are subjected to fluid shear. We present a theoretical framework within which we calculate the fraction of detached cells as a function of shear stress for fast ramps as well as for the decay in the fraction of detached cells at fixed shear stress as a function of time. Using HEK and 3T3 fibroblast cells as experimental model systems, we extract characteristic force scales for cell adhesion as well as characteristic detachment times. We estimate force-scales of ~ 500 *pN* associated to a single focal contact, and characteristic time-scales of 190 ≤ *τ* ≤ 350s representing cell-spread-area dependent mean first passage times to the detached state at intermediate values of the shear stress. Variations in adhesion across cell types are especially prominent when cell detachment is probed by applying a time-varying shear stress. These methods can be applied to characterizing changes in cell adhesion in a variety of contexts, including metastasis.

## 1. Introduction

Cells that adhere to substrates provide a convenient model system to explore the magnitude and distribution of the forces that cells exert on the extracellular matrix (ECM). The adhesions that cells form with the ECM are mediated principally by a class of adhesion molecules called integrins. In their bound state with ECM receptors, a number of other proteins bind to integrins, forming a complex structure called a focal adhesion. Such focal adhesions (FAs) contain a large number of transmembrane and cytosolic proteins with force-sensitive components, such as talin and vinculin [1, 2, 3], making them sensitive mechanosensors [4, 5]. Focal adhesions are also dynamic entities, which respond to external stimuli via mechano-chemical feedback processes. By linking the actin cytoskeleton and the ECM, focal adhesions can function both as force and biochemical sensors, linking mechanosensing with downstream signalling cascades that enable cells to modify their behaviour in response to perturbations.

Cell migration, proliferation [6], differentiation [7, 8] and cellular metastasis [9, 10] are all cellular processes that involve the modulation of cell adhesion[11, 12]. Changes in cell adhesion are also seen in certain disease states. Adhesion complexes are known to be modified in cancer cells [13] which have been shown to have a higher spread area, to apply large traction forces on their surroundings [14, 15], and to proliferate faster [16]. Cells which detach from tumors and metastasize appear to have enhanced focal adhesion dynamics [17], raising questions of how the modification of cell adhesion in disease might lead to impaired force sensing, with downstream consequences for tumour spreading. Such observations also suggest that quantifying variations in the adhesion of cells cultured on substrates might provide interesting insights into this problem.

Investigating cell adhesion at a quantitative level is difficult because adhesions are dynamic and heterogeneous in their sizes and spatial distribution. As a result, depending on the length scale of interest and the questions at hand, several distinct approaches have been used to probe cell-substrate interactions. At a molecular level, atomic force microscopy has been used to study the attachment and detachment dynamics of single bonds [18]. Micron sized beads coated with extracellular matrix proteins can be attached to cells and pulled using an AFM or a magnetic tweezers [19]. At the single cell level, cell detachment can be studied using micropipettes [20, 21]. Finally, population-averaged cell-detachment kinetics can be studied using an external fluid shear stress applied by means of a spinning disc [22, 23] or by using microfluidic flow channels [24]. Centrifugation forces have also been used to study cell detachment [25]. Population studies conducted using fluid flow stress or centrifugation are of special importance when comparing differences in gross adhesiveness between cell types. Such population-based methods provide powerful tools to compare normal cells with cells with specific knock-down or knock-out modifications to the focal assembly. From a bio-medical point of view these techniques have the potential to investigate changes in adhesion brought about by metastasis [26]. Quantifying cell-substrate adhesion is also essential for the design of compatible synthetic implants [25]. Some of the techniques mentioned above have been recently reviewed [27].

If the anomalous behavior exhibited by cells with compromised adhesion are associated to extreme values of a distribution [28], it is unlikely that assays based on single cells will be sensitive to them. On the other hand, population studies, if based on a sufficiently large population, are likely to accommodate a range of varied cell adhesions, including these extreme ones. Additionally, collective phenomena such as wound healing and metastasis are examples where population studies are more appropriate than single cell studies. Another advantage of population-based studies is that they connect more effectively with theoretical models based on positing a distribution of adhesion strengths and numbers of focal adhesions. Data testing these models is most efficiently accumulated through population studies rather than through single-cell studies.

A controllable way of measuring the distributions of adhesion strength is to subject adhered cells to fluid shear[29]. The most common ways of applying the fluid shear to study the cell detachment kinetics are by means of a spinning disc apparatus [22, 30] or through the use of microfluidic flow channels [24]. However, extracting cell adhesion parameters from such measurements requires the development of theoretical models. A theoretical model for the effects of a shear flow on adherent cells must account for the following: Forces acting on the cell arising from the shear flow must be computed accurately. These will depend on cell geometry as well as its modulation by the applied force. Next, the effects of these forces as they act on cell-substrate attachment points must be computed. A probabilistic model for the distribution of attachment points is essential for the understanding of results from population-based studies of cell detachment under an applied force. Finally, the stochastic dynamics of the attachment and de-attachment of individual focal adhesions under external forces must be accounted for. Despite the complexity of this problem, one might hope that simplified models, incorporating what one believes to be essential ingredients, might provide useful insights [31].

This paper presents a shear-device-based cell detachment assay coupled to a theoretical model that describes how the size distribution of the adherent region in specific cell types should influence cell-substrate adhesion strengths. We use our shear device to generate data for the force-induced de-adhesion of 3T3 fibroblast and HEK293T cells from fibronectin-coated substrates, both at varying shear stress as well as at constant shear stress as a function of time. To understand these results, we develop a theoretical model for the modulation of cell adhesion under an applied force, illustrating how adhesion parameters can be extracted from the experiments. We use a load-sharing assumption for the action of these forces on a prescribed number of adhesion points and a generalization, based on a stochastic version of the Bell model, for how de-adhesion occurs at fixed shear stress.

Our model assumes that the forces required to detach cells from their substrates should scale simply with the size of the adhered region, a specific hypothesis that we use to motivate and derive analytic expressions that describe how cells de-adhere upon the application of a force. We consider specific forms for the distribution of cell sizes, relating them to the distribution of attachment points. We use these results to obtain analytic results for the fraction of cells that remain adhered, upon a fast ramp of the applied shear rate, as well as a formula for the decay of the number of adhered cells with time at an intermediate value of the shear rate. These analytic forms derive directly from the assumptions made for the distribution of cell sizes, as well as for the distribution of the number of attachment points at given spread cell radius, but can be generalised to arbitrary distributions. Our fits to these functional forms agree well with the experimental data, demonstrating both the usefulness of this device in probing cell adhesion as well as its usefulness, once coupled to the model description, in extracting numerical values of relevant biophysical parameters from population-based assays.

## 2. Materials & Method

### 2.1. Cell Culture

3T3 fibroblast and HEK293T cells were grown on plastic petri dishes coated with fibronectin. The coating was done by exposing the surface to 30 *µ*g/ml fibronectin (Sigma Aldrich) in HBSS for 30 min. The cells were grown in DMEM containing 10% FBS and 1% PSG to about 50% confluency so that they do not touch each other. All reagents were purchased from Invitrogen. After plating, the cells were incubated overnight in 5% CO_2_ at 37 °C before experiments.

### 2.2. Imaging and Analysis

In order to quantify the number of cells detached as a function of time and shear stress under fluid shear, we counted labelled nuclei. The cell nuclei were labelled using Hoechst H33342 (Molecular Probes) at 1:1000 ratio of the 10 mg/ml slock solution. The fluorescence imaging was done using a motorized Zeiss AxioObserver-Z1 microscope, an Axiocam CCD camera, and the AxioVision 4.8 software, all supplied by Zeiss. The counting of labelled nuclei was done using in-house MATLAB code that involved counting of bright objects of a specified size range. Another in-house code was written in MATLAB to measure cell areas. Both the codes were written using the in-built Matlab function “regionprop”.

### 2.3. Device

Our custom-built shear device uses the standard cone and plate geometry to apply controlled fluid shear stress to adherent cells. This custom device is compact and inexpensive and can be mounted on any standard inverted microscope and imaging can be performed in phase contrast, fluorescence, confocal mode etc. For enabling high resolution imaging. More details of the electronics and image of the device are provided in 1 and Supplementary Fig. S1. A computer hard disk motor removed from a SATA, 3.5”, 7200 rpm PC hard disk drives a vertically mounted cone plate at a desired rpm. This precision motor is almost perfectly wobble free and allows for precision coupling of the cone plate with excellent matching of the motor and cone axes. The 1° cone plate and shaft are fabricated out of a single aluminium piece and the cone surface is polished for smoothness. The rpm of the three phase, out-runner, brushless motor is controlled via an electronic speed controller (ESC) (HobbyWing 80020591) which in turn is controlled using pulse width modulation signals from a function generator (Agilent Technologies, 33220A). The actual rpm depends on the load on the motor, which varies with the experimental condition. To measure the actual rpm, the shaft of the cone is painted black except for a thin reflective strip. An encoder unit consisting of an IR LED and a sensor (TCRT5000 reflective sensor module) measures the reflected light pulses and records the rpm on a computer using an Arduino Uno USB I/O board.

The petri dish containing adherent cells is placed below the cone plate and its temperature is controlled using heating foils (Minco ribbon heaters) attached to the bottom aluminium base plate and controlled using an calibrated Pt-100 RTD and a temperature controller (CT325 Minco). Three micrometer levelling screws (Thorlabs Inc., KS1, Precision Kinematic Mirror Mount) on the motor mount allows for setting the rotation axis of the cone normal to the surface containing the cells. This alignment is done iteratively by measuring the gap between the bottom flat surface and the top cone plate at three points equidistant from the cone apex. The distance and gap measurements are performed with a resolution better than a micrometer using the motorized XYZ units (with encoders of the Zeiss AxioObserver Z1 microscope), and a long distance 40*×* objective. The final gap is set by first lowering the cone using the vertical micrometer stage (Thorlabs Inc., MT1, Travel Translation Stage, least count 10*µ*m) till its apex gently presses on the cells below and this point is set as zero. The cone plate is then raised to the desired working distance. The imaging for cell counting was done using a 10× objective.

## 3. Results and Discussion

### Cell detachment assays

We use two protocols to study the detachment kinetics of adhered cells. In the first, a constant fluid shear stress is applied and the number of cells detached was recorded as a function of time. In the second, the imposed shear stress is increased in steps with a fixed waiting time between steps, with the number of remaining cells counted at the end of each wait time. Our results, obtained using fibroblasts and HEK as model systems, are shown in Fig. 2. The constant shear used in the first protocol and the waiting time in the second protocol were decided from prior trials such that the constant shear stress is set to equal the threshold stress seen in Fig. 2b where 50% of the cells are detached. The wait time is chosen to be of the order of the characteristic detachment time obtained in constant stress experiments as the one shown in Fig. 2a. Similar detachment curves have been reported earlier but the quantification we report here is novel[32, 33, 34]. The number of attached cells as a function of increasing shear clearly shows an initial plateau where cells do not detach over the times spent at each stress value, since the shear stress is much less than the threshold stress defined by the mid-point of the curves in Fig. 2b.

**Figure 1.**
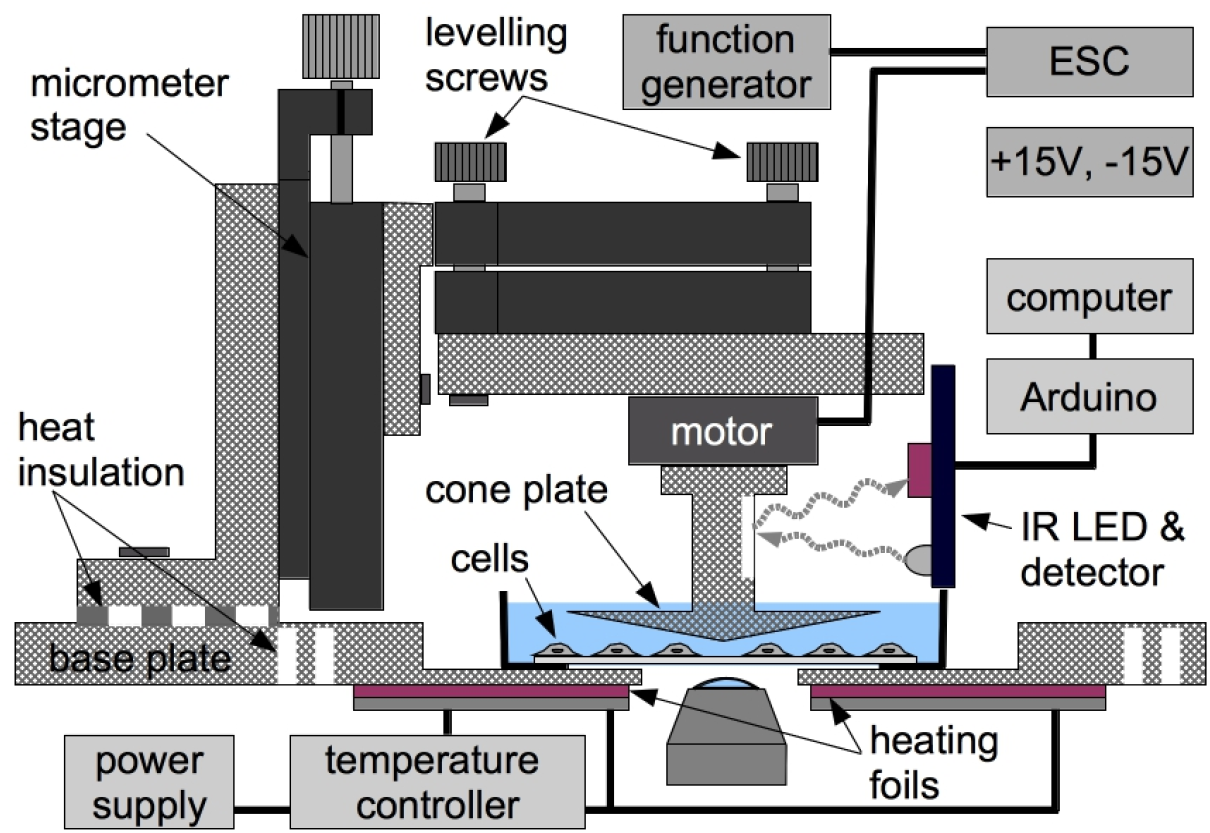
A schematic of the shear device fabricated using a hard disk motor. The rpm of the motor is controlled using an electronic speed controller and a function generator. The rpm is measured using an IR LED and sensor coupled to a computer via an Arduino board. The temperature of the petri dish containing cells is controlled using an electrical temperature controller. The cone is aligned with respect to the petri dish containing the cell monolayer using three micrometer levelling screws. The cone to plate gap is adjusted using a vertical micrometer translation stage. The gap and tilt measurements were performed using encoders on a motorized microscope. Imaging is done in fluorescence mode.

**Figure 2.**
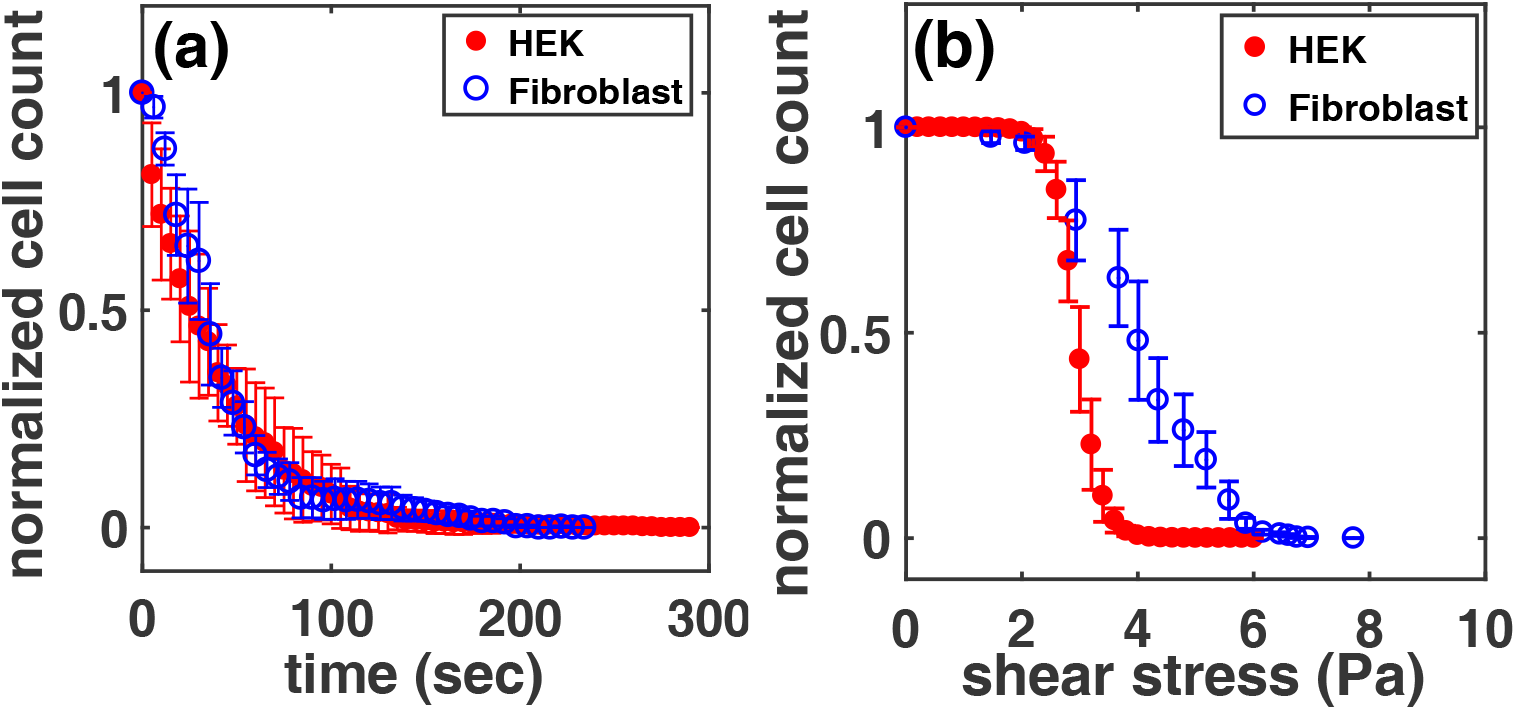
a) Detachment kinetics at a constant fluid shear stress. The value of the shear stress was taken to be that corresponding to the value at which 50% of cells had detached in each case, as extracted from (b), (b) Detachment curves for HEK and fibroblast cell as a function of shear stress with a waiting time of 1 min at every shear value. The curves shown in both (a) and (b) represent an average detachment trend, taken over 4 independent runs. The starting cell count in both cases, for both the cell types, was around 100. Error bars are *±* standard deviation (SD).

The number of cells that remain adhered as a function of time at a constant shear stress appears to decay largely as an exponential, although there is a small but significant deviation from this form in the tail region for the 3T3 fibroblast cells. This is possibly due to a small population of well-spread cells of much larger-than-average area in the case of the fibroblast cells, as can be seen in Fig. 3. Note that the 3T3 cells have an average spread area of 1800 *µ*m^2^ whereas the HEK cells we used have a significantly lower spread area of 369 *µ*m^2^ as shown in Fig. 3. This figure also exhibits numerical fits to the areal distribution for both HEK and fibroblasts in terms of a log-normal distribution, as discussed further below. A theoretical explanation of the dependence of detachment curves upon the distribution of spread areas of cells is provided in the following section.

**Figure 3.**
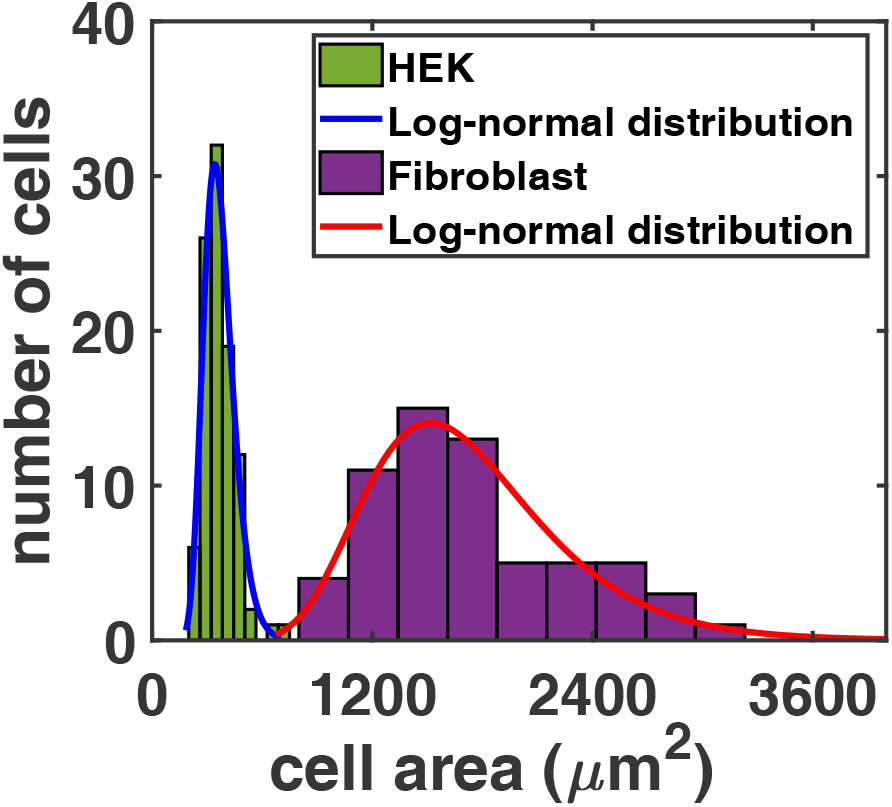
Cell spread area distributions measured for 3T3 fibroblasts and HEK cells as indicated in the figure legend. These indicate that the broader detachment curves in the case of the fibroblasts likely originates in the overall broader distribution of cell sizes. The solid curves in both cases are fits to a log-normal distribution. The parameters that fit the distribution in each case are: *µ* = 5.88, *s* = 0.22 (HEK) and *µ* = 7.41, *s* = 0.299 (Fibroblasts) where *µ, s* are the mean and standard deviation of the associated normal distribution. The total numbers of cells taken for the distributions are 62 for 3T3 fibroblasts and 100 for HEK cells.

### Theoretical Model

To estimate the scale of forces exerted on adherent cells in a shear flow, we use results from Price, Pozrikidis and other workers[35, 36, 37, 38, 39, 40]. Consider shear stresses acting across a solid hemisphere attached to a flat substrate. We assume that the flow is governed by the linear Stokes equation, and is constrained by incompressibility[41, 42]. For such a shear flow, with the velocity in the x-direction and the gradient in the z-direction, with shear rate *k* and fluid dynamical viscosity *η* and thus wall-shear stress *σ* = *ηk*, the total force exerted on the hemisphere is 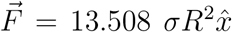 where *R* the cell radius[36]. We ignore effects due to the torque exerted by the flow, confinement effects which would correct the coefficient above, viscoelastic response of the cell, as well as possible feedback between the flow and cell orientation and shape, although some of these effects can be significant [43, 44, 43, 45, 46, 47]. Taking *R* ~ 20*µm* (*R*_*HEK*_ ≃ 11*µm, R*_3*T*3_ ≃ 24*µm*) and values of *σ* of around 3.5*Pa* (*σ*_*c*_ ≃ 3*Pa* for HEK cells and *σ*_*c*_ ≃ 4*Pa* for 3T3 cells) required for detachment, we get a force-scale of *~* 19 *nN*, in the right range as compared to typical experimental values for the total force required for cell detachment[24, 41, 48, 31, 49, 50]. Estimating the number of adhesion complexes to be around 40, this implies that each adhesion complex exerts approximately 475 *pN* of force on the substrate, a figure in the appropriate range *vis a vis*. experiments[48]. Modelling adherent cultured cells as hemispheres attached to a rigid substrate ignores some important aspects of their geometry. In interphase, such cells are typically well-spread on substrates, with their maximal heights *h* typically smaller than the spread radii *R*. The maximal height is controlled by the placement and size of the nucleus and the structure of the F-actin cortical layer [51]. Pozrikidis provides numerical solutions for a family of protuberances involving sections of oblate spheroids and semi-spheroids, finding that the scaling of total force with the contact area remains quadratic, although it is modified by a (geometry-related) pre-factor which is in the range of about (1-14) [35, 52]. One way of accounting for cell shape changes resulting from shear would be to allow this numerical factor to vary with shear stress [42]. Other calculations treat the cell-fluid interface not as a solid-liquid interface as we do here but as an interface between two fluids of very different viscosities [38]. Provided this viscosity contrast is large, our results should continue to hold.

Here, we have simply equated the forces exerted on the cell by the flow to the cell adhesion forces, ignoring the dynamics of attachment and detachment of adhesion molecules. However, cell adhesion is a collective phenomenon. A mean-field approach to understanding the unbinding and rebinding of adhesion molecules under force *F* follows from the work of Bell, who considered an adhesion cluster as a set of *N* molecules capable of binding as well as detaching from a substrate [53]. At time *t*, *N* (*t*) are bound while the remaining *N − N* (*t*) adhesion molecules are unbound. Each bond breaks with a rate *k*_*off*_ and can rebind with a rate *k*_*on*_. The incorporation of a force promotes unbinding via *k*_*off*_ = *k*_0_ exp (*Fx*_*b*_/*k*_*B*_*TN*(*t*)) = *k*_0_ exp (*F*/*F*_*b*_) where *x*_*b*_ is a typical bond length scale, *T* is the temperature, *k*_*B*_ the Boltzmann constant and *F*_*b*_ is a molecular scale force. From the discussion above we can assume that *F*_*b*_ ≃ 0.5*nN*. It is conventional to assume that the load is equally shared by the bound adhesion molecules. These approximations then lead to the mean-field equation for *N*(*t*),

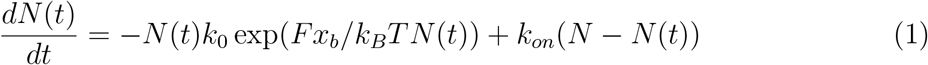

This is simplified by introducing a dimensionless time *τ* = *k*_0_*t*, rescaling force as *f* = *F/F*_*b*_, scaling the rebinding rate so that *γ* = *k*_*on*_/*k*_0_ and by assuming that a constant force is applied and shared equally between all adhesion molecules. In steady state, this yields *N* (*τ*) exp(*f/N* (*τ*)) = *γ*(*N − N* (*τ*)), an equation which predicts a saddle node bifurcation when *f*_*c*_ = *Np* ln(*γ/e*), where the product logarithm is defined as *p* ln(*a*) from the solution of *x* exp *x* = *a*. This implies that adhesion is stable up to a critical force *f*_*c*_ in mean-field theory[54, 55, 31].

Such a mean-field description ignores fluctuations. Stochasticity can be incorporated in terms of a one-step master equation for the quantity *p*_*i*_, where *p*_*i*_ is the probability of having *i* adhesion molecules bound at time *t*[54, 31, 56, 55, 57]. Such an equation incorporates both loss and gain terms and can be written as

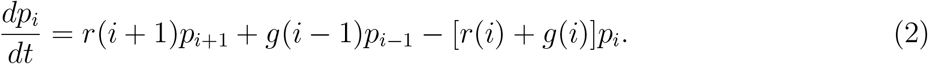

With rates appropriate for the Bell model with load sharing *r*(*i*) = *i* exp(*f/i*) a*nd g*(*i*) = *γ*(*N − i*), the mean first passage time *T* can then be obtained from a result of Van Kampen[54, 31]. This result yields a logarithmic dependence of *T* on the force for small forces and an exponential dependence at large forces, with the cross-over defining a characteristic (dimensionless) force value *f*_*c*_. Note that adhesion involving an finite number of adhesion molecules, each with a characteristic unbinding force, must ultimately be unstable, although the characteristic time for detachment can be large, depending on *N*. This treatment also suggests two experimentally relevant regimes, the first in which the mean number of attachment points is instantly equilibrated as the shear stress is ramped in steps but any further fluctuations are ignored. This is essentially a short-time or fast-loading approximation. A second limit is one in which we wait at each imposed shear stress value for times comparable to the first passage time *T*, in which case we must account for the dynamics of detachment in detail.

We draw intuition from kinetic Monte Carlo methods that incorporate stochasticity to compute the fraction of bound cells at time *t* following the application of the flow, in the short-time limit described above. We simulate directly the stochastic process defined by the equations above using the Gillespie method [58, 59, 60]. We determine the average fraction of cells which are adhered at a particular level of shear, given that they are observed *up to* a given time *t*_0_, as measured from the initiation of the shear flow. Our calculations first assume a fixed initial number of adhesion molecules (50 *−* 100), simulated using Eq. 2. Our choice of rates follows that of the Bell model. Our results are shown in Fig. 4(a). where we have varied *N*, the mean number of attachment points at zero force. Note that there is an essentially sharp transition between a state in which the number of attachment points varies from a value close to *N* to zero. The force value at which this transition occurs is subject to hysteresis between different runs, but the values cluster about a mean detachment force that scales linearly with *N*, as in the simple Bell model. Fig. 4(b) shows that as the waiting time at each force value increases, the threshold value decreases, leaving intact the step jump of this fraction at a waiting-time-dependent critical force. The systematic decrease of the threshold with increasing waiting time also reflects the fact that cell detachment in the presence of stochastic fluctuations involves a barrier crossing process. We model the experimental data by using the result that the critical force for the abrupt detachment of a cell with *N* equivalent attachment points increases linearly with *N*, *F*_*c*_(*N*) = *αN*.

**Figure 4.**
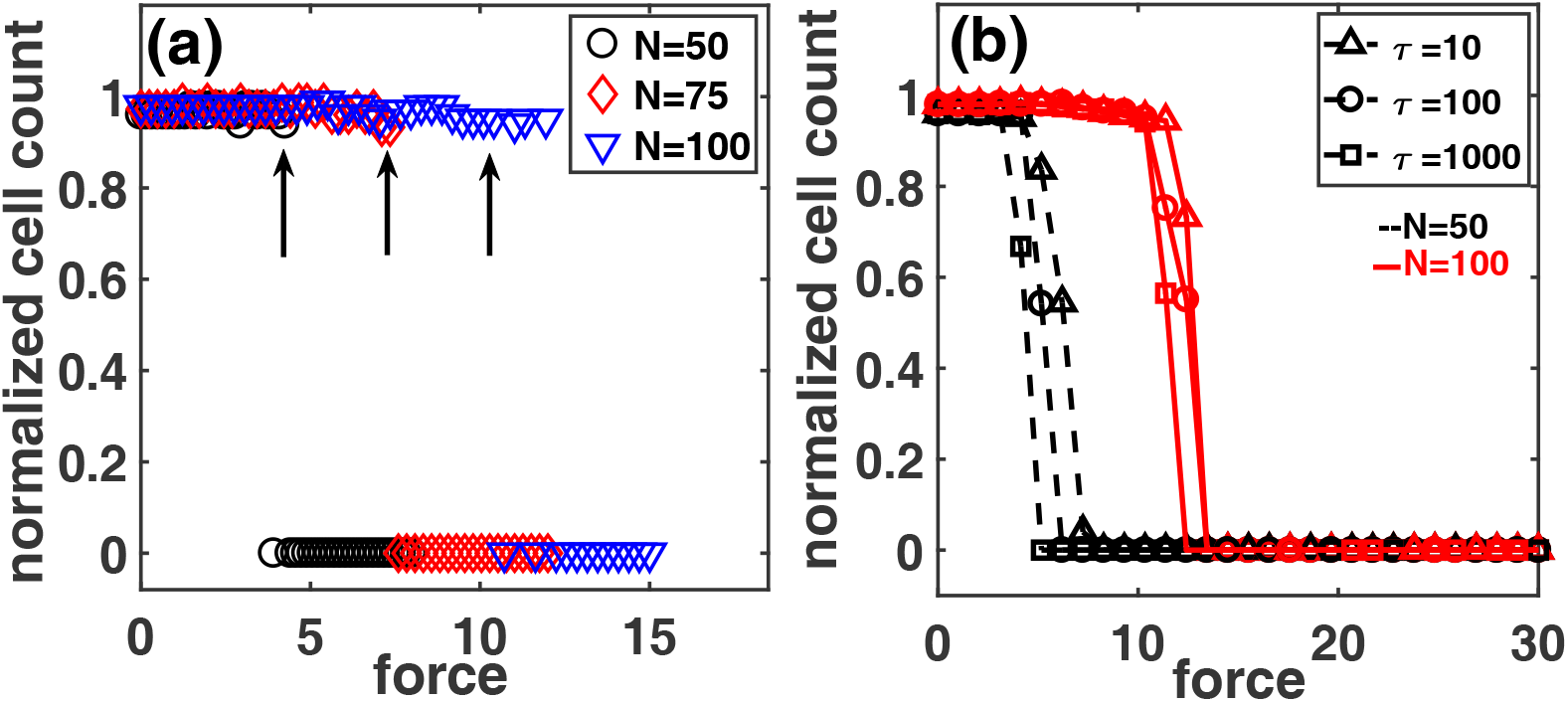
(a) Fraction of adhered cells as a function of force acting on the adhered cell via a shear flow, as computed using kinetic Monte Carlo methods as appropriate to the Bell model. This is shown for three values of *N*, the steady state number of attachment points at small times in the absence of a force as well as for a number of different runs. Note the sharp drop in this fraction near a critical force *f*_*c*_ (vertical arrows) which scales linearly with the number of initial attachment points; the location of this drop can vary across runs as seen in the hysteretic behaviour displayed, as is characteristic of first-order transition behaviour, but its location clusters about the average value indicated by the arrow. All cells are assumed to be identical in size. (b) Fraction of adhered cells as a function of applied force. Each set of curves for a given N (given color) are for different waiting time (different symbols) at each force value varied over a logarithmic scale. As the waiting time is increased, the threshold value decreases, while the essentially sharp decrease of this fraction at the waiting time and N-dependent threshold value remains a feature of the data..

We can relate the applied shear stress *σ* to the total force *F* experienced by each cell, which will, in general, depend on the shape of the cell and the characteristic length-scales over which the flow is perturbed. Our basic question relates to the fraction of adhered cells Φ(*σ*) that are observed to survive when the shear stress is increased to *σ* from zero. Φ(*σ*) will depend on the history of the shear, both if we assume that detached cells are lost via flow outside the imaging region as well as if we take into account the stochasticity in the detachment and attachment of focal adhesions.

For concreteness, we will consider the case where the shear stress is ramped up fast from zero, essentially ensuring that only those cells which are absolutely unstable to detachment are removed, returning later to the case in which we wait at each stress value for a defined time, as in the experiments. The derivative

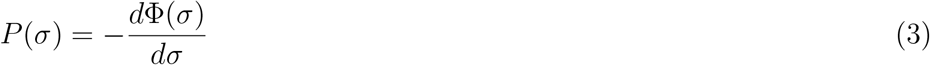

represents the fraction of cells that detach between *σ* and *σ* + *dσ*. Thus, we have

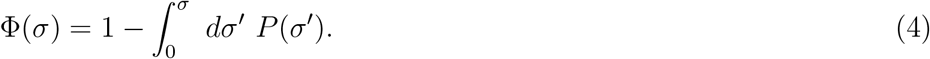

We wish to calculate *P* (*σ*) for a set of adhered cells with the quantities *R* and *N* given by a joint distribution *P* (*N, R*). This is given by

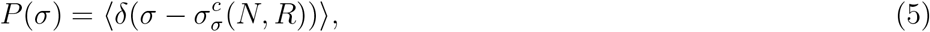

where the averages, denoted by 〈·〉, are over the probability distribution *P* (*N, R*). The shear stress *σ* is related to the force exerted on the cell by the flow and the critical value of the shear stress is denoted by 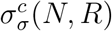. Thus,

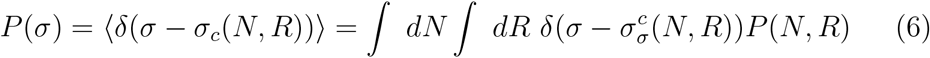

Now, we can decompose this joint probability in terms of the conditional probabilities

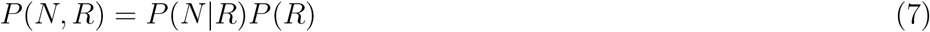

We will make the approximation that the conditional probability of having *N* attachment points is slaved to the radius *R* and has a particularly simple form,

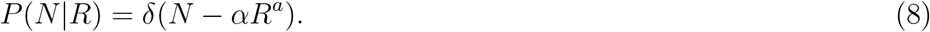

The distribution of cell sizes then determines *P* (*σ*) completely. Then, the probability distribution of critical depinning forces given a probability distribution of *R* is

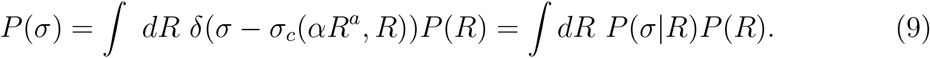

The shear force experienced by our model cell is 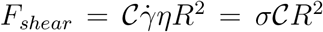, with 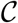 a geometric factor reflecting the aspect ratio of the adhered cell, *η* is the viscosity, *σ* the wall stress and *R* the radius of the circular section in contact with the substrate. This force is opposed by forces from the FA’s: *F*_*adhesions*_ = *Nf* = *αR*^*a*^*f*, where we have assumed that the number *N* of focal adhesions is directly proportional to *R* raised to an appropriate power and the magnitude and dimensions of *α* defined accordingly. If FA’s are distributed largely along the perimeter, then *a* = 1, which is the limit we consider. Equating these, we obtain 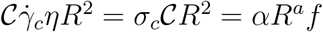, which provides an estimate for the critical shear stress defined through

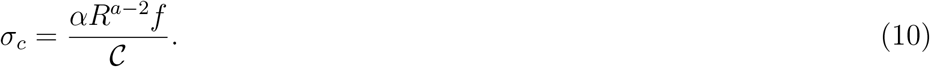

When *a* = 1, corresponding to FA’s concentrated along the perimeter of the cell, we have

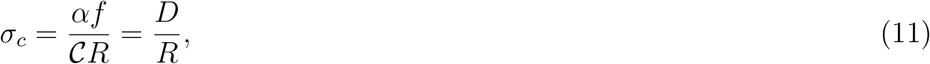

where 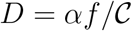

Assuming that the distribution of spread cell sizes is log-normal[61], we have

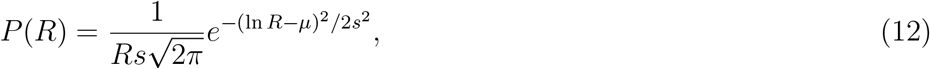

where *µ* and *s* are the location and scale parameters of the associated normal distribution. Given

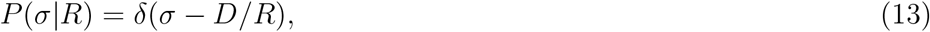

we obtain

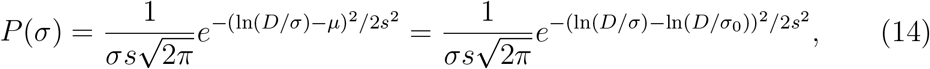

with

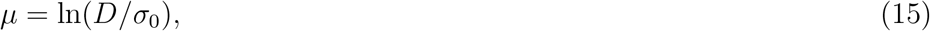

which we simplify as

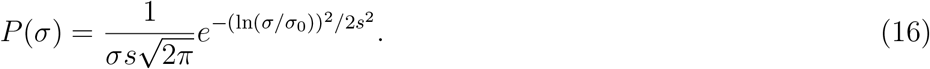

Now we have

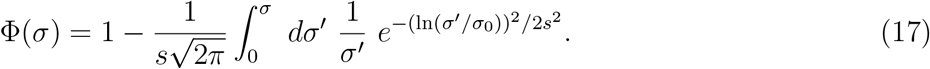

Using the definition of the error function

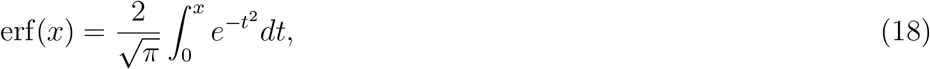

this finally yields our central result

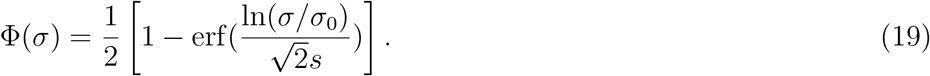

When *σ* = *σ*_0_, Φ(*σ*) = 1*/*2, defining the critical force *σ*_0_ as the point where half of the cells have detached upon a fast ramp.

This expression, Eqn. 19, contains fit parameters *σ*_0_ and *s*. The first of these, *σ*_0_, can be directly inferred from data, since it is just the value at which half of the cells have detached. We optimise the value of *s* to obtain the best fits. In Fig. 5 we show the experimental data of Fig 2 against our prediction of Eqn. 19. For the data shown in Fig. 5(a), these parameters are: *σ*_0_ = 2.91, 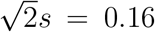 (HEK) and in Fig. 5(b) *σ*_0_ = 3.85, 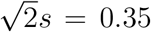 (Fibroblasts). The quantity *s* is proportional to the width of the distribution of the number of attachment points across cells, which we assume is equivalent to the distribution of cell radii. We have also implicitly assumed that all adhesion points are alike.

**Figure 5.**
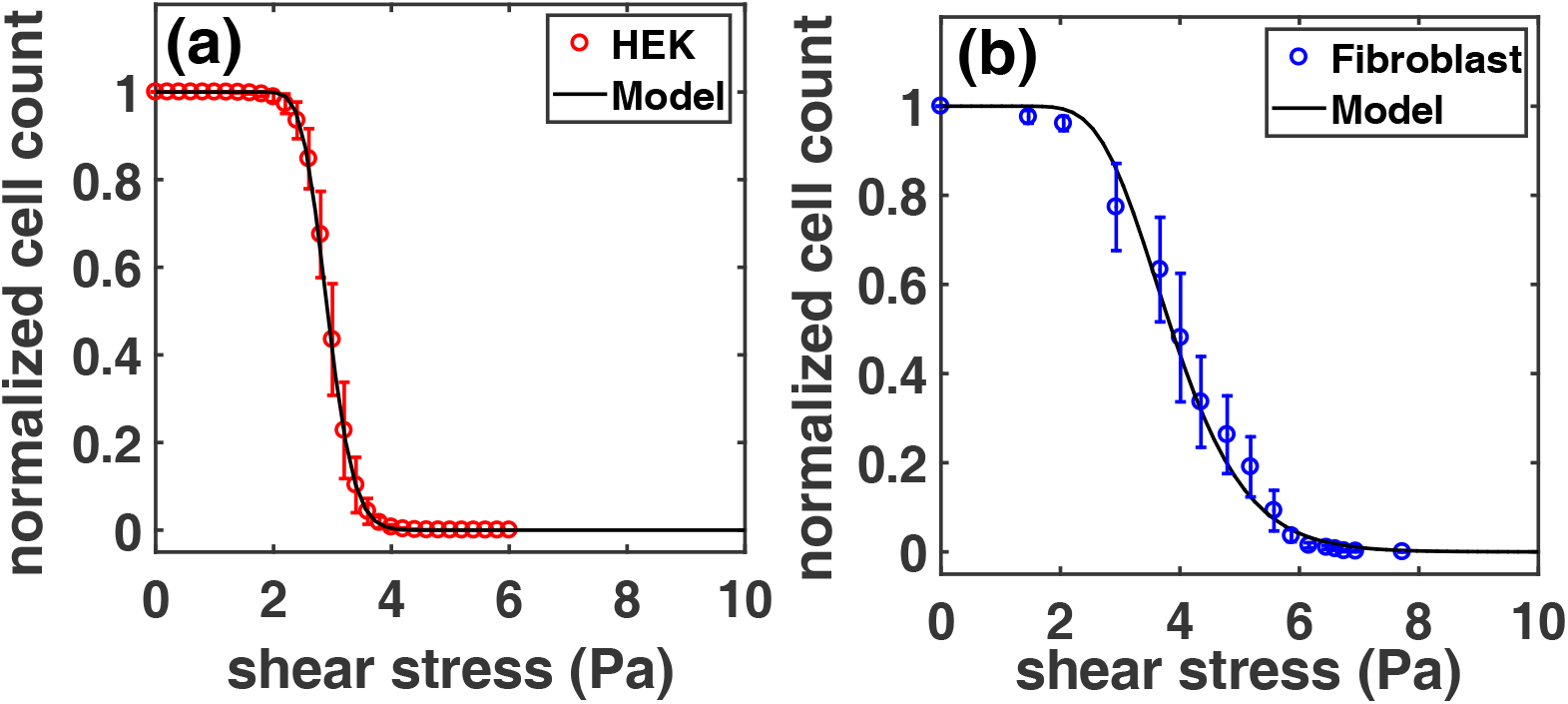
(a) Plot of the experimental data for HEK cells shown in Fig 2 against the model prediction of Eq. 19 with the values of *σ*_0_ and *s* obtained from a best-fit analysis and (b) Plot of the experimental data for fibroblast cells as shown in Fig 2 against the model prediction of Eq. 19 with the values of *σ*_0_ and *s* obtained from a best-fit analysis.

To understand the behaviour in the limit that we spend a sufficiently long time at each force value, we must take into account the dynamics of the adhesion molecules, in particular the fact that fluctuations can carry an otherwise stable cluster of attachment points to a regime where adhesion becomes unstable. It is simplest to assume that *σ*_0_ as a function of time decays as an exponential with a characteristic time scale *τ*, since this is consistent with the absence of a force dependence of the first passage time in this limit. (More complex relaxations can easily be accounted for). We thus have,

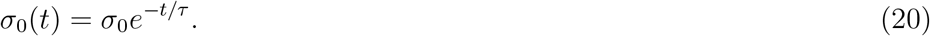

This then yields,

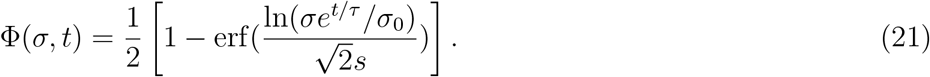

One further simplification is obtained when *σ* = *σ*_0_, *i.e*. we examine relaxation at the value of the shear stress where 50% of cells are unstable to detachment upon a fast ramp. In this limit, and normalizing to the number of cells present at *t* = 0, we have,

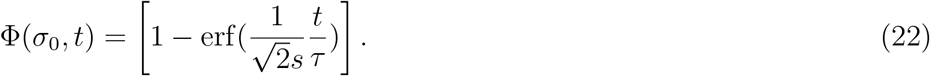

In Fig. 6 we show the data for both HEK and fibroblasts, where the fit form is obtained by using the values of 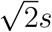 from the prior fits for the separate cases of HEK and fibroblasts and the value of *τ* is changed till an optimum fit is obtained. In this manner we obtain *τ* = 190*s* for fibroblasts and *τ* = 350*s* for HEK cells. These time-scales are perhaps most accurately as averaged first passage times to the detached state, where the average is taken over a distribution of number of attachment points across cells of different spread sizes as well as over a large number of stochastic realisations.

**Figure 6.**
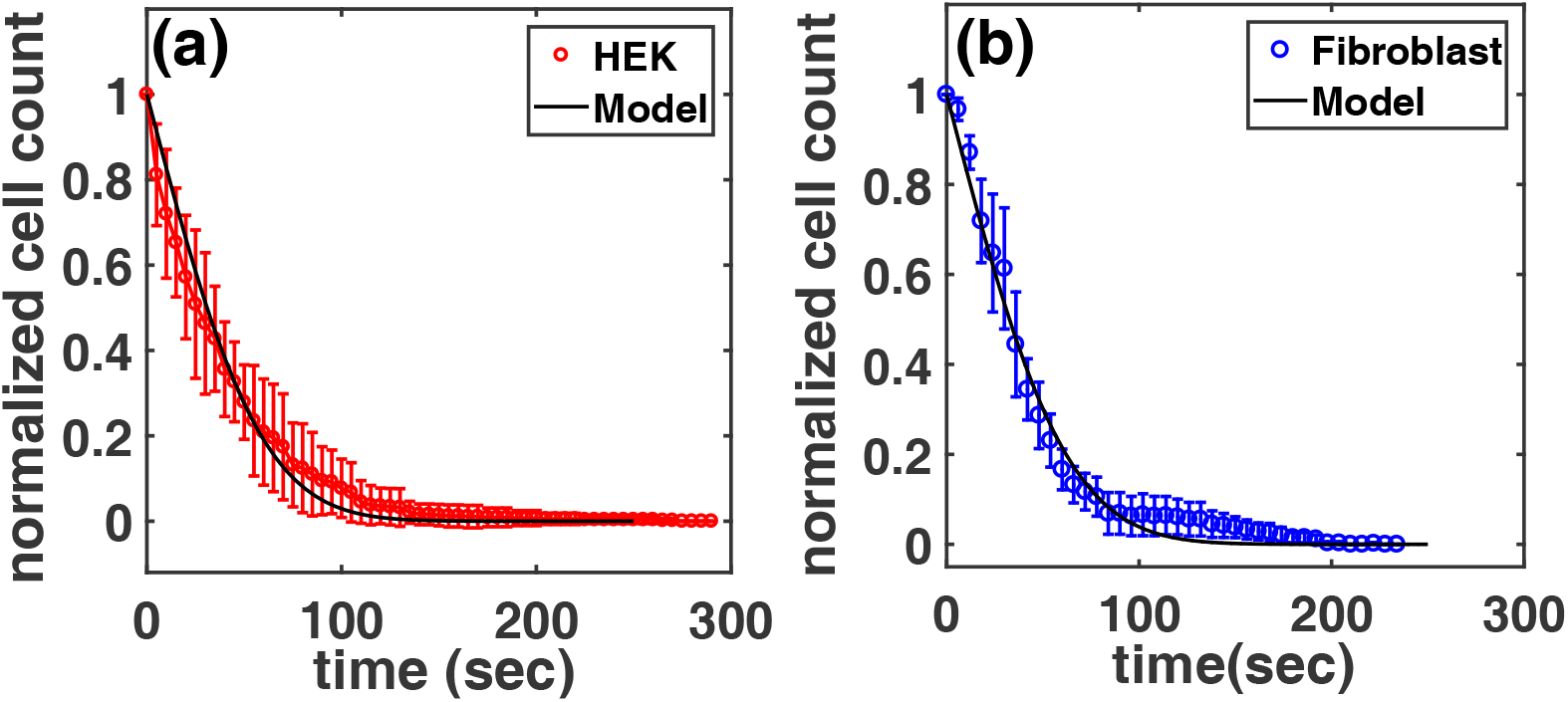
(a) Plot of the fitting function 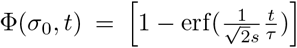 against the experimental data for detachment as a function of time at fixed stress *σ*_0_ for HEK cells, (b) Plot of the fitting function 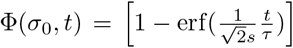 against the experimental data for detachment as a function of time at fixed stress *σ*0 for fibroblasts. In both cases, we take the values of *σ*_0_ to be the midpoint value of the detachment curve, i.e. the value of the stress at which, on a fast ramp, 50% of cells are observed to detach. We take the values of *s* from fits to the detachment curve, leaving *τ* as the only undetermined parameter which we optimise to obtain the best fit.

In principle we might have expected, given the theoretical development, that the value of *s* obtained from fitting the distribution of cell spread areas to a log-normal should be related to the one that is used to fit the detachment data which uses the fit of the cell radii to a similar log-normal form: If *A* ~ Lognormal(*µ, s*^2^) then 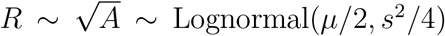, where *s*^2^ denotes the variance here. In practice, there is a difference between these values: Assuming *P* (*R*) consistent with the fits to the area distribution in Fig. 3, we would predict standard deviations for the distribution of radii to have values 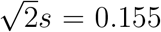 for HEK cells and 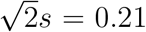 for fibroblast cells. From Fig. 5, our best fit values are 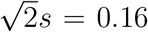 (HEK) and 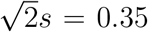 (fibroblasts). The relative closeness of fit values for HEK cells likely reflects the quality of the fit, as can be seen from Fig. 3. The fit to a log-normal distribution for the spread area of fibroblasts is certainly inferior, possibly accounting for the discrepancy.

One way to generalize the approach above is to allow for different sub-populations of cells, governed by independent log-normal distributions of cell spread areas. We could have, for example, cell spread areas distributed according to two independent distributions as 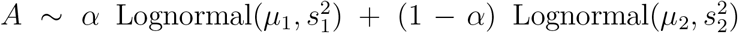, with 0 ≤ α ≤ 1 denoting the relative weights of the two (normalized) distributions used to fit the experimental data. The distribution of spread cell radii would then follow 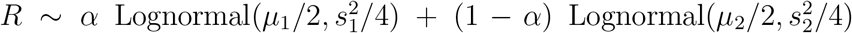 in this case. The related expression for Φ(*σ*) follows in a straightforward manner as derived above, with the parameters entering this expression related to *s*_1_ and *s*_2_ and the weights of the constituent distributions for the cell radius. These expressions can be generalized easily to further sub-populations.

We checked whether the fits shown in Fig. 5 (b) could be improved upon by fitting the spread area histograms for fibroblasts shown in Fig. 3 to more complex distributions in place of a single log-normal. Our results, shown in Supplementary Information (Figs. S2 and S3), indicate that the detachment curve can be fit to reasonable accuracy (Fig. S3) if we assume that the spread area data arises from summing over three appropriately weighted independent log-normal distributions (Fig. S2), with parameter values indicated in the figure captions. However, the numbers of cells, 62 for the 3T3 fibroblasts, is clearly too small for us to infer with any confidence that this reflects the underling biology.

This difference could also originate in the other approximations we made: First, we ignored heterogeneities in the number of detachment points *N*, assuming that *N* scales simply with the radius and is uniquely prescribed, once *R* is supplied. Second, we idealised all cells as characterized by only their spread area, via a single quantity *R*, and assumed that cell-spread areas were distributed according to a log-normal distribution. Third, we ignored variations in the strength of different focal adhesions. Given these somewhat stringent approximations, the fact of a discrepancy between the theoretically predicted values extracted from the cell-size distribution data and the ‘best-fit’ values of *s* extracted from the detachment curves is unsurprising.

Finally, a functional form similar to the one we derive (Eq. 19) for the detachment curve has been proposed earlier, in Ref. [34]. However, there it was simply used as a fitting formula, motivated by the expectation that detachment thresholds at a particular shear rate should arise from the product of several “independent and random factors”. If these individual factors are drawn from Gaussian distributions, the distribution of their product takes a log-normal form and the appropriate cumulative distribution has the form of Eq. 19. Here, in contrast, we show how such a formula can be explicitly derived, given simple assumptions whose structure we motivate here. We relate parameters that are treated simply as fit parameters in Ref. [34] to measurable quantities such as the variance in the spread cell area and the strength of individual focal adhesions. The advantage with such an explicit treatment is that it can be generalized, by relaxing each of the assumptions we mention above. In particular, the calculated detachment curve would have a different form if we assumed a form other than the log-normal for the distribution of spread cell areas.

## Conclusions

This paper has described population-based measurements of cell detachment from substrates upon the application of a fluid shear stress. Such measurements relied on the development and benchmarking of a compact and inexpensive cell shearing device, while the interpretation of the results used a model which integrated a number of different approaches to this problem. Our device is made of inexpensive components, such as a high precision computer hard-disk motor from a discarded disk controlled by an electronic speed controller and a Arduino board. The micrometer adjustment screws and the heating elements can also be implemented easily using ready made and inexpensive items. The use of a 30mm coverglass as the bottom surface of the petri allows for in-situ high resolution imaging. As it is extremely compact, the device can be easily mounted on any standard inverted microscope. It can thus be used in combination with different types of fluorescence microscopy techniques, such as confocal and total internal reflection (TIRF) microscopy.

Although spinning disk devices have been used in the past to study cell detachment under shear stress there are a few drawbacks which prevent them from being more widely employed. The use of a commercial rheometer as in Derks et al. [62] is attractive due to the ease of use and the range of accessible protocols but such devices are prohibitively expensive if used primarily for cell detachment studies. Shear devices can also be technically challenging to fabricate [63]. Several of the reported custom devices are unsuitable for in-situ visualization of cell detachment dynamics and require post-analysis by removing the cell culture plate [23, 64]. Microfluidic flow channels such as described in Decave et al., Steward et al., Couzon et al., and Lu et al. [65, 66, 24, 67] are compatible with microscopy but achieving high enough flow rates to study cell detachment is difficult. Such devices are often prone to disastrous leaks when used with thin coverglasses (for high resolution) on expensive microscopes. Besides this, even with recirculation, flow technique requires a large amounts of culture medium for long term experiments. This becomes an issue especially when using expensive biochemical reagents (drugs) that perturb cell adhesion properties. A comprehensive review of the different techniques for the measurement of cell adhesion, describing their relative advantages and applications, can be found in Ref. [27]. Our device overcomes some of the difficulties mentioned above. In addition, a significant aspect of the present study is that we have performed de-adhesion experiments in two complementary modes: constant shear stress and time-varying shear stress.

The theoretical model we preset here integrates elements from a stochastic description of the formation and breakage of individual adhesion molecules with a cell-scale description of the forces experienced by cells when the surrounding fluid is sheared. Our analysis indicates that even if we assume that all focal adhesions are identical, accounting for variations in cell size and any accompanying variation in the number of attachment points across cells could account quantitatively for the shapes of the detachment curves, as seen in the quality of the fits to the formulae we derive. This suggests that our methodology can be used to distinguish delicate differences in cell adhesion across different cell lines or cells with different modifications. For the two cell lines we study in this paper, these differences can be argued to arise directly from the distribution of cell sizes which is different in both cases. This distribution influences the predicted detachment curves, since it determines the width of the detachment curve. Specifically, we show that the time varying shear stress mode is better at discerning differences in the adhesion of HEK and 3T3 cells. Since, in particular, cancerous cells are known to differ from normal cells in their adhesion strengths, the device can potentially be used to distinguish between such cells using a population-based measurement [26].

Our theory accommodates an arbitrary distribution of cell sizes, although for simplicity we confined ourselves to calculations for the log-normal distribution. We have suggested that, in the simplest case of a homogeneous cell population, a single set of parameters *σ*_0_ and *s*, should fit both the constant shear stress and time-varying shear stress data. In particular, the quantity *s* can be interpreted as the width of the distribution of the number of attachment points, which is equivalent, apart from a dimensional multiplicative factor, to the distribution of spread cell radii in the simple model described here. Given the generality of the approach described here, it is easy to accommodate more complex joint distributions *P* (*R, N*) as well as to incorporate more detailed biophysical modelling of variations in adhesion strengths.

A further improvement of the setup would involve the use of micro-patterned substrates to restrict cells to well defined sizes and shapes. In this case, we can test our assumptions of simple linear relationships between the number of adhesion points, the radius corresponding to the spread area of the cell and inhomogeneities of the force scale of different attachment points. Working with synchronised cells will reduce one source of uncertainty in the data. The possibility of “catch-bonds”, in which bond dissociation lifetimes increase with an applied tensile force, can be incorporated into the theoretical framework reported here. Another central question relates to the description of the feedback between the flow and cell-scale bio-physical properties, a many-scale problem that links molecular scale force-sensing and signaling to changes in the shape of the cell and its orientation with respect to the flow. Incorporating these modifications to the experiment and analysis should improve our understanding of, as well as our ability to quantitatively describe, the biophysical underpinnings of cell adhesion phenomena.

## Acknowledgments

We gratefully acknowledge funding from the PRISM project of the Institute of Mathematical Sciences, Chennai. We are particularly grateful to Namrata Gundiah for useful discussions and to Rahul Siddharthan and Jean-Fran¸cois Joanny for productive conversations. We thank Nidhi Pashine for her contribution to the development of the shear device.

